# Cytochrome OmcS is not essential for long-range electron transport in *Geobacter sulfurreducens* strain KN400

**DOI:** 10.1101/2020.07.22.214791

**Authors:** David J. F. Walker, Yang Li, David Meier, Samantha Pinches, Dawn E. Holmes, Jessica A. Smith

## Abstract

The multi-heme *c*-type cytochrome OmcS, is one of the central components for extracellular electron transport in *Geobacter sulfurreducens* strain DL-1, but its role in other microbes, including other strains of *G. sulfurreducens* is currently a matter of debate. Therefore, we investigated the function of OmcS in *G. sulfurreducens* strain KN400, which is even more effective in extracellular electron transfer than strain DL-1. We found that deleting *omcS* from strain KN400 did not negatively impact the rate of Fe(III) oxide reduction and did not affect the strain’s ability to accept electrons via direct interspecies electron transfer. The OmcS-deficient strain also continued to produce conductive filaments, consistent with the concept that electrically conductive pili are the primary conduit for long-range electron transfer in *G. sulfurreducens* and closely related species. These findings, coupled with the lack of OmcS homologs in most other microbes capable of extracellular electron transfer, suggest that OmcS is not a common critical component for extracellular electron transfer.

## Introduction

*Geobacter sulfurreducens* strain DL-1 is one of the most widely studied microorganisms capable of extracellular electron transfer. Studies have indicated that extracellular electron transfer by DL-1 is complex and involves a multitude of components, most-notably the electrically conductive pili (e-pili) (1, 2) and numerous outer cell-surface *c*-type cytochromes (3). One of the most important and thoroughly studied *c*-type cytochromes involved in strain DL-1 extracellular electron transfer is OmcS. OmcS, is a six-heme *c*-type cytochrome (4), associated with the outer cell surface and the e-pili of DL-1 (5). Strain DL-1 requires OmcS for growth on insoluble Fe(III) oxide, but not soluble Fe(III) citrate (6). OmcS is also essential for direct interspecies electron transfer (DIET) in co-cultures of DL-1 and *Geobacter metallireducens* (7), but not for long-range electron transport through current-producing biofilms of strain DL-1 (8-10).

Under some conditions OmcS assembles into µm-long filaments (11-12). Based on these findings it was suggested that the formation of OmcS filaments “explains the remarkable capacity of soil bacteria to transport electrons to remote electron acceptors for respiration and energy sharing” (12). However, OmcS homologs are found in only a few soil bacteria that are capable of extracellular electron exchange (2, 13). Furthermore, many bacteria capable of extracellular electron exchange do not produce any outer-surface cytochromes (14-16). OmcS homologs are not even prevalent within the genus *Geobacter* (13). For example, *G. metallireducens*, a close relative of *G. sulfurreducens*, does not contain any OmcS family proteins, yet its rate of Fe(III) oxide reduction is significantly faster than *G. sulfurreducens* (17). *Geobacter uraniireducens*, one of the few microbes that has an OmcS family protein (13), lacks phenotypes associated with long-range electron transport through conductive filaments. For example, *G. uraniireducens* is incapable of participating in DIET and cannot produce high current densities on anodes (17). *G. uraniireducens* transfers electrons to Fe(III) oxides with a soluble electron shuttle, not through filament-mediated direct electron transport (18).

In order to further evaluate whether OmcS is an important component for extracellular electron transfer in any microbes other than *G. sulfurreducens* strain DL-1, we studied *G. sulfurreducens* strain KN400, a strain that is more effective in extracellular electron transfer than the DL-1 strain (19, 20). We found that deleting the gene for OmcS had no impact on extracellular electron transfer or the production of electrically conductive filaments, suggesting a limited significance for OmcS beyond *G. sulfurreducens* strain DL-1.

## Results and Discussion

### An OmcS-deficient mutant of *G. sulfurreducens* effectively reduces Fe(III) oxide

A strain of *G. sulfurreducens* KN400 in which *omcS* was deleted reduced Fe(III) oxide just as fast as the wild-type strain during the active phase of Fe(III) oxide reduction (Fig. 1). In the initial transfer from fumarate medium, the lag period prior to rapid Fe(III) oxide reduction was longer in the OmcS-deficient strain (12 days) than for the wild-type strain (5 days), but it is difficult to attribute physiological significance to this longer initial lag period. For example, the construction of the mutant required selection and recovery on medium with the soluble electron acceptor fumarate, a growth condition that can deadapt cells for growth via extracellular electron transfer (10). After several transfers, the lag phase in the OmcS*-*deficient mutant was comparable to the wild-type and both strains continued to reduce Fe(III) oxide at similar rates (Fig. 1b). These results demonstrated that OmcS was not required for effective Fe(III) oxide reduction.

**Figure 1.**
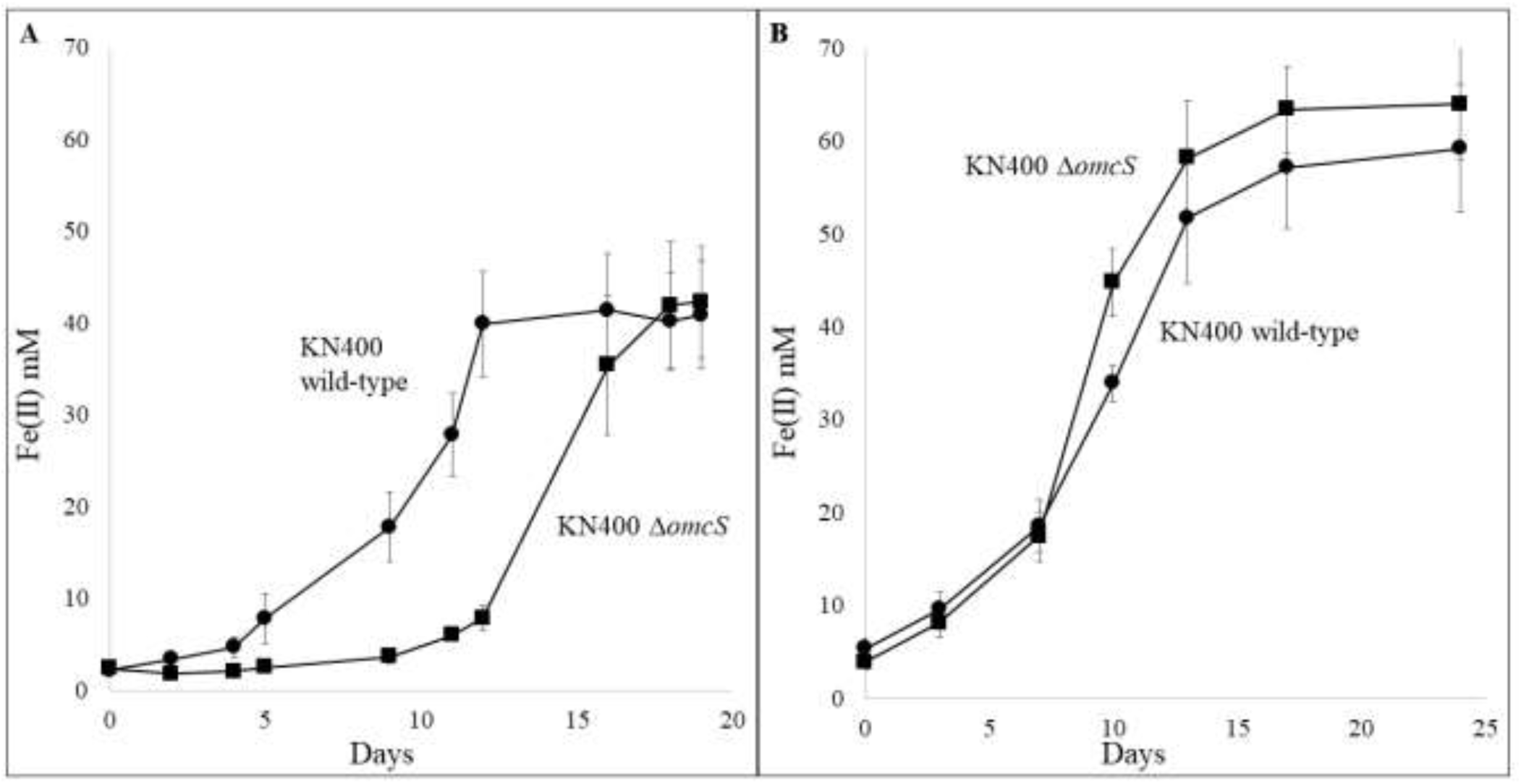
(a) Fe(II) production from Fe(III) oxide reduction over time in the KN400 wild-type and OmcS-deficient strains after the first transfer into Fe(III) oxide medium from fumarate medium and (b) Fe(II) production from Fe(III) oxide reduction over time in the KN400 wild-type and KN400 OmcS-deficient strain after five consecutive transfers in Fe(III) oxide medium.

### Deletion of *omcS* does not increase expression of the electron shuttle PgcA

A previous study of a *pilA*-deletion mutant in KN400 demonstrated that after hundreds of days a strain could be adapted to effectively reduce Fe(III) oxide (21). This adaptation was attributed to a mutation that increased expression of the soluble extracellular *c*-type cytochrome PgcA (KN400_1784), which can function as an extracellular electron shuttle to facilitate electron transfer to Fe(III) oxides (21, 22). In order to evaluate the possibility of increased PgcA expression in the OmcS-deficient mutant, outer cell surface proteins were sheared from Fe(III) oxide-grown cells, and combined with cell free filtrate. The proteins were analyzed on SDS-polyacrylamide gels stained for heme (Fig. 2a). PgcA, a 41 kDa protein, was not abundant in either the wild-type or OmcS-deficient strain. In fact, transcript abundance for the gene for PgcA was lower in the OmcS-deletion strain than the wild-type strain (Fig. 2b).

**Figure 2.**
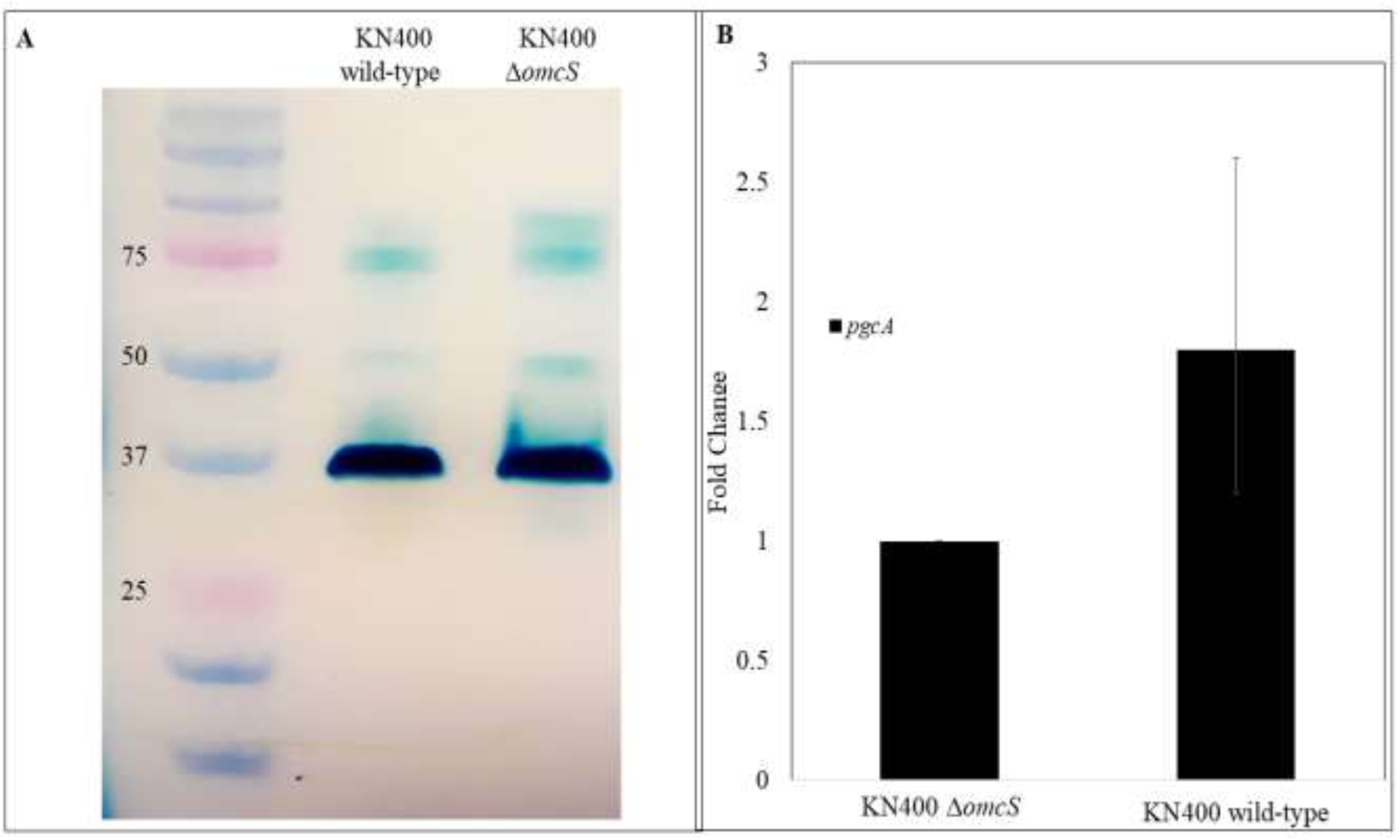
Lack of increased cytochrome expression in the OmcS*-*deficient mutant. (a) *c*-type cytochrome content of outer surface protein and cell-free filtrates from the wild-type and OmcS-deficient cultures of KN400 grown with Fe(III) oxide as the electron acceptor. Proteins were separated by SDS-PAGE and stained for heme. The intensely stained band below the 37 kDa marker is likely OmcZ, which has previously been shown to be highly expressed in strains of KN400 adapted for Fe(III) oxide reduction (21). (b) Transcript abundance of *pgcA* in the wild-type and OmcS-deficient cultures of KN400 grown with Fe(III) oxide as the electron acceptor. Relative expression ratios were determined by the 2^-^_Δ Δ_^Ct^ method (23), using the housekeeping gene *proC* for data normalization. Triplicate technical and biological replicates were done for all samples.

### Impact of OmcS deletion in DIET co-cultures

Previous studies have shown that OmcS is also required for DIET in co-cultures of *G. sulfurreducens* strain DL-1 and *G. metallireducens* (7). In fact, long-term growth of strain DL-1 via DIET selected for a strain with a mutation in a regulatory gene which resulted in increased expression of OmcS (7). Therefore, the OmcS-deficient strain of KN400 was tested in DIET co-cultures with wild-type *G. metallireducens*. Similar to the Fe(III) oxide reduction results, deletion of OmcS in KN400 had no effect on DIET even in the initial co-cultures (Fig. 3).

**Figure 3.**
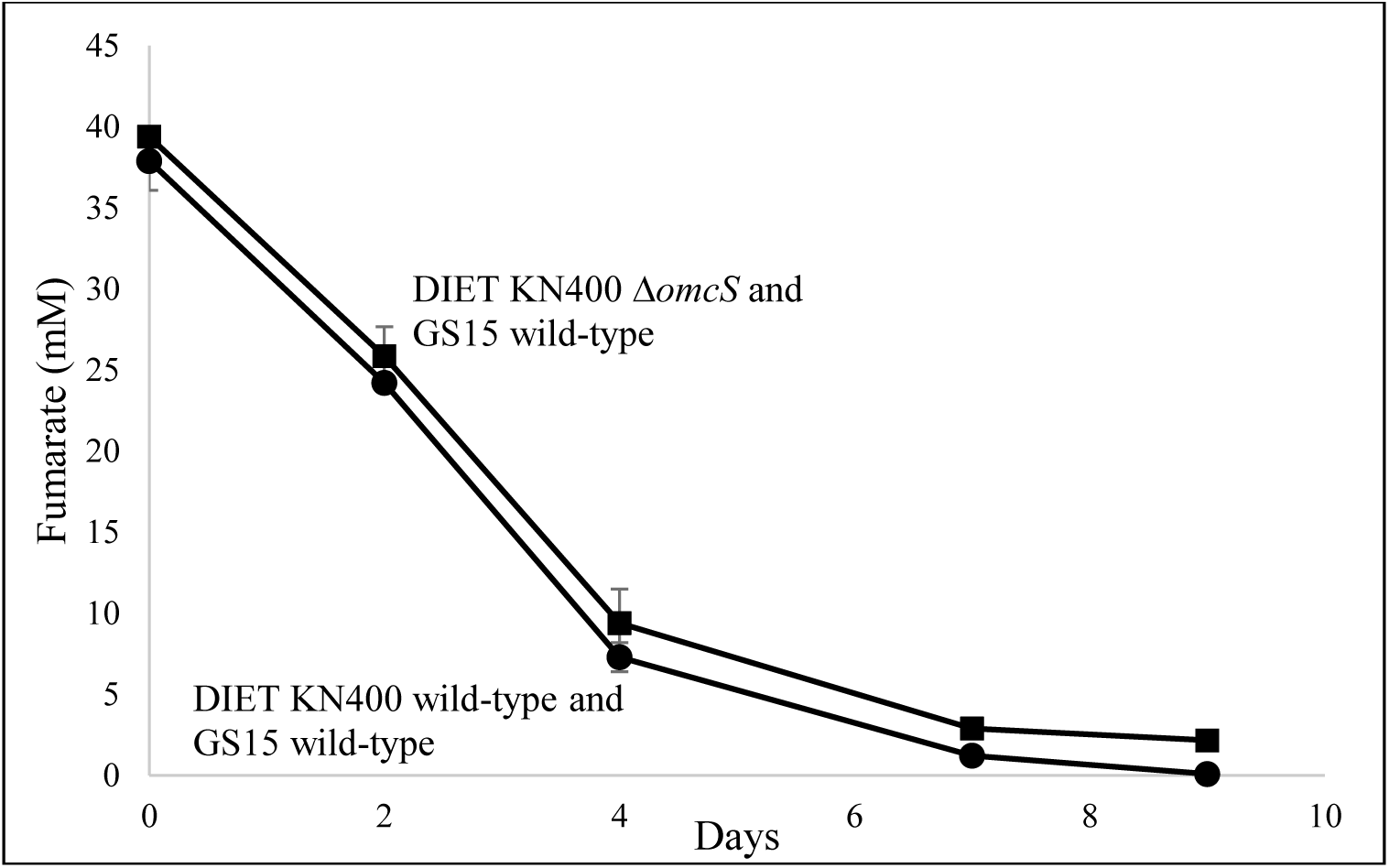
Metabolism of DIET co-cultures initiated with the wild-type or OmcS-deficient strain of *G. sulfurreducens* KN400 and wild-type *G. metallireducens* GS15.

### The filaments of the OmcS deletion strain are still conductive

As previously reviewed (1, 2), many studies have suggested that the primary conductive filaments produced by *G. sulfurreducens* strain DL-1 are e-pili. To determine whether the OmcS-deficient strain of *G. sulfurreducens* KN400 continued to produce conductive filaments, filaments were sheared from the cells and their conductivity evaluated with 4-probe solid-state I-V measurements. The electrical conductance of filaments sheared from KN400 Δ*omcS* cells was comparable to the previously reported (16) conductance of *G. sulfurreducens* strain DL-1 e-pili networks, and much higher than filaments from strain Aro-5, which expresses poorly conductive pili (Fig. 4).

**Figure 4.**
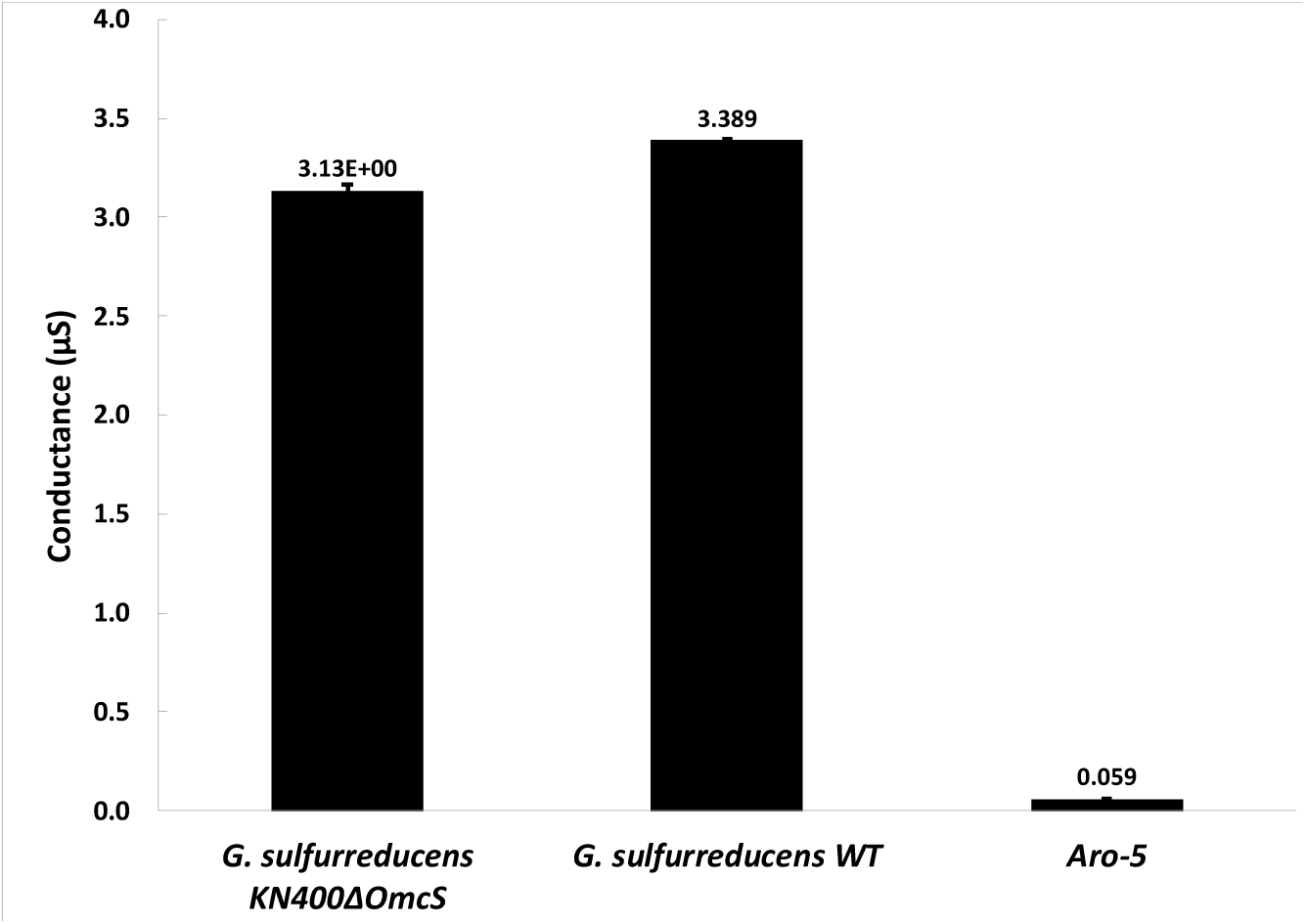
Four-probe conductance (mean ± standard deviation; n=9) of filament networks purified from *G. sulfurreducens* KN400 Δ*omcS*, compared with previously reported conductances (16) of filaments harvested from wild-type *G. sulfurreducens* strain DL-1, and strain Aro-5.

### Implications

The results presented here demonstrate that *G. sulfurreducens* strain KN400 does not require OmcS for effective extracellular electron transfer or production of electrically conductive filaments. These results suggest that the importance of OmcS in extracellular electron transfer is limited, even amongst the *sulfurreducens* species of *Geobacter*.

It has been suggested (12) that filaments comprised of OmcS, rather than e-pili, are the predominant conductive filaments expressed by *G. sulfurreducens*, but multiple lines of evidence have refuted this hypothesis (2). For example, when a short peptide tag was included in a synthetic gene for the pilin monomer, all filaments observed emanating from the cell contained the peptide tag (24). In addition, the conductivity of filaments harvested from *G. sulfurreducens* can be increased or decreased by modifying the abundance of aromatic amino acids in the pilin monomer (18, 25-29), a modification that would not be expected to influence the conductivity of OmcS filaments. Also, construction of strains that produce an abundance of OmcS, but express synthetic pilins that yield poorly conductive pili, are defective in long-range electron transfer (18, 25-26, 29). The findings reported here, and in another recent study (30), show that deleting the gene for OmcS has no impact on the conductivity of the filaments harvested from *G. sulfurreducens*, further demonstrating that OmcS filaments cannot be the primary conductive protein nanowires expressed by *G. sulfurreducens*. Thus, the actual role of OmcS in *G. sulfurreducens* strain DL-1 warrants further study.

## Materials and Methods

### Bacterial strains and growth conditions

*Geobacter sulfurreducens* strain KN400 (20), *G. metallireducens* strain GS15, and the Δ*omcS* strain of *G. sulfurreducens* DL-1 (6) were obtained from our laboratory culture collection. The *G. sulfurreducens* strains were routinely cultured under strict anaerobic conditions (80% N_2_ and 20% CO_2_) with 10 mM acetate provided as the sole electron donor and either 100 mM Fe(III) oxide or fumarate (40 mM) provided as the sole terminal electron acceptor (31, 32). Cultures were incubated at 30 °C. The medium composition per liter of deionized water with either Fe(III) oxide or fumarate as the sole electron acceptor was prepared as follows: 0.42 g KH_2_PO_4_, 0.22 g K_2_HPO_4_, 0.2 g NH_4_Cl, 0.38 g KCl, 0.36 g NaCl, 0.04 g CaCl_2_ 2H_2_0, 0.1 g MgSO_4_ 7H_2_O, 1.8 g NaHCO_3_, 0.5 g Na_2_CO_3_, 1.0 ml of 100 mM Na_2_SeO_4_, 10.0 ml of DL vitamin solution listed in (33), and 10.0 ml of NB trace mineral solution listed in (31). For genetic manipulation to create KN400 Δ*omcS*, cells were grown in either liquid or agar-solidified acetate-fumarate medium (31), and grown in an anaerobic chamber with an N_2_-CO_2_-H_2_ (83%, 10%, and 7%, respectively) atmosphere at a temperature of 30 °C.

All pure culture strains of *G. metallireducens* were regularly transferred into FC medium (32) with 20 mM ethanol provided as the sole electron donor and 56 mM Fe(III)-citrate as the sole electron acceptor. DIET co-cultures were initiated with equal amounts of both organisms in anaerobic pressure tubes containing 10 mL of NBF medium, with 10 mM ethanol provided as the sole electron donor and 40 mM fumarate as the electron acceptor.

### KN400 *omcS*-deletion mutant construction

To construct the KN400 *omcS*-deletion mutant, primers OmcS_fwd (GGTCGTGATGCTCGATCCGGAAG) and OmcS_rev (GGTTGGCGGTAAGGAGGTGCCG) were used to amplify the mutated region from the DL-1 *omcS*-deletion strain. The PCR product was purified using a QIAquick gel extraction kit (Qiagen, Valencia, CA) according to the manufacturer’s instructions, and verified by Sanger sequencing. Electrocompetent *G. sulfurreducens* strain KN400 cells were prepared from cultures maintained on acetate-fumarate medium, and all manipulations were carried out on ice in an anaerobic chamber with a N_2_-CO_2_-H_2_ (83%, 10%, and 7%, respectively) atmosphere at a temperature of 30°C. 200 ml of cells were harvested (when they reached an optical density at 600 nm of 0.2) by centrifugation for 8 minutes and 4300 x g at 4°C. Cells were washed twice with electroporation buffer (1 mM HEPES pH 7.0, 1 mM MgCl_2_, and 175 mM sucrose). Final cell pellets were resuspended in electroporation buffer to achieve a final concentration of 10^11^ cells/ml. 100 ng of the purified PCR product was electroporated into KN400 cells at 14.7 kV/cm for 6 ms (31). The cells were then inoculated into fumarate-acetate medium and allowed to recover for 8 hours at 30°C, prior to plating on fumarate-acetate agar with spectinomycin (25 μg/ml). Spectinomycin was added for selection purposes only. Replacement of wild-type alleles by mutant alleles was verified by PCR and Sanger sequencing.

### Cell-free filtrate SDS-PAGE and protein identification

Outer cell surface and cell-free proteins in the medium were collected by centrifugation at 17700 x g for 20 minutes from 50 ml cultures of KN400 wild-type and KN400 Δ*omcS* cells grown with Fe(III) oxide provided as the electron acceptor when Fe(II) concentrations reached approximately 40 mM. Loosely bound outer cell surface proteins from one hundred milliliters of culture were sheared with a Waring blender operated at room temperature at low speed for 2 minutes (6, 34). Blended samples were then centrifuged at 10,000 × g for 20 min to remove cells and insoluble Fe(III) oxide, and supernatant was collected and concentrated with an Amicon Ultra-15 centrifugal filter unit (Millipore, Billerica, MA). Proteins were quantified with a Micro BCA protein assay kit (Thermo Scientific, Rockford, IL) using the manufacturer’s instructions. Equal amounts of protein (5 μg) were separated by SDS-PAGE in glycine-buffered 12.5% polyacrylamide gels. Heme groups were identified using the heme stain to detect peroxidase activity with H_2_O_2_ and 3,3’,5,5’-tetramethylbenzidine (TMBZ) (35).

### RT-qPCR

Total RNA was extracted from triplicate cultures during exponential growth in acetate (10 mM)-Fe(III) oxide (100 mM) using the HG method (36). For this extraction, 50 ml of cells were pelleted by centrifugation at 3000 x g for 20 minutes at 4°C and immediately resuspended in 10 ml of preheated (65°C) HG buffer (100 mM Tris-HCl pH 8, 100 mM NaCl, 10 mM EDTA, 2.5% β-mercaptoethanol, 1% sodium dodecyl sulfate, 2% Plant RNA Isolation Aid (Ambion, Woodward, TX, USA), 5 mM ascorbic acid, 0.6 mg/ml Proteinase K, and 5 mg/ml lysozyme). The mixture was incubated at 65°C for 10 minutes, and then 2 μl Superase-In (Ambion) and 0.025 mM CaCl_2_ were added. After centrifugation for 10 minutes at 16,100 x g, the supernatant was transferred to new 2 ml screw capped tubes. 50 μl of Plant RNA Isolation Aid (Ambion), 4 μl of 5 mg/ml linear acrylamide (Ambion), 600 μl of preheated (65°C) acidic phenol pH 4.5 (Ambion), and 400 μl of chloroform-isoamyl alcohol (24:1; Sigma, St. Louis, MO, USA) were added, and rotated for 10 minutes. The samples were then centrifuged for 5 minutes at 16100 x g, at which point the aqueous layer was removed and transferred to a new tube containing 100 μl of 5 M ammonium acetate (Ambion), 20 μl of 5 mg/ml glycogen (Ambion), and 1 ml of cold isopropanol. The samples were left to precipitate at -30°C for 1 hour.

After precipitation, the nucleic acids were pelleted by centrifugation at 16100 x g for 30 minutes. The pellet was washed twice with 70% ethanol, dried, and resuspended in sterile diethylpyrocarbonate-treated (DEPC) water (Ambion). The RNA was purified with a RNeasy MiniElute cleanup kit (Qiagen) according to the manufacturer’s instructions and treated with Turbo DNA-free DNase (Ambion). The RNA samples were tested for genomic DNA (gDNA) contamination by PCR amplification of the 16S rRNA gene. The concentration and quality of the RNA samples were determined with a NanoDrop ND-1000 spectrophotometer (NanoDrop Technologies, Wilmington, DE) and by agarose gel electrophoresis. All RNA samples had A260/A280 ratios of 1.8 to 2.0, indicating high purity. cDNA was generated with a TransPlex whole-transcriptome amplification kit (Sigma-Aldrich, St. Louis, MO) according to the manufacturer’s instructions.

Primer pairs with amplicon sizes of 100 to 200 bp were designed for the *G. sulfurreducens pgcA* (pgcAF: ACGGGAATAACTGTCAGGCA and pgcAR: CGTGAGGGTTGAAAGGAACC) gene. Expression of this gene was normalized with expression of *proC* (proCF: ACCGATGACGATCTGTTCTTT and proCR: ATGAGCTTTTCCTCCACCAC), a housekeeping gene constitutively expressed in *Geobacter* species (37). Relative levels of expression of the studied genes were calculated by the 2^−ΔΔ*CT*^ threshold cycle (CT) method (23).

Power SYBR green PCR master mix (Applied Biosystems, Foster City, CA) and an ABI 7500 real-time PCR system were used to amplify and to quantify the PCR products. Each reaction mixture consisted of forward and reverse primers at a final concentration of 200 nM, 5 ng of cDNA or gDNA, and 12.5 μl of Power SYBR green PCR master mix (Applied Biosystems).

### Analytical techniques

Organic acids were monitored with high performance liquid chromatography (HPLC) as previously described (38).

Fe(II) concentrations were measured with the ferrozine assay (39). For this assay, samples of Fe(III) oxide cultures were dissolved at a 1:10 ratio in 0.5 N HCl for 2 hours at room temperature. Next, 0.1 mL of sample extract was added to 5 ml of ferrozine (1 g/L in 50 mM HEPES buffer at pH 7) and mixed. Concentrations of Fe(II) were determined by measuring absorbance at 562 nm in a split-beam dual-detector spectrophotometer (Spectronic Genosys 2; Thermo Electron Corp., Mountain View, CA).

### Purification of e-pili

The e-pili were harvested from 100 ml of stationary phase cultures of *G. sulfurreducens* KN400 Δ*omcS* grown with acetate as the electron donor and fumarate as the electron acceptor at 30° C. The e-pili were purified as previously described (40). Briefly, the e-pili were separated from cells using a blender in 150 mM ethanolamine (pH 10.5) and the cells were then separated via centrifugation. The e-pili-containing supernatant was subjected to two rounds of ammonium sulfate precipitation, and the purified e-pili were resuspended in ethanolamine buffer and stored at 4°C. Ethanolamine was then removed from the e-pili samples with dialysis tubing (Slide-A-Lyzer, 2KDa, ThermoFisher) suspended in deionized water. Concentrations of purified protein were measured with the Pierce nano bicinchoninic acid assay (Thermo Scientific), as previously described (40) and samples were normalized to 500 μg/μl.

### Four-probe solid state thin film e-pili conductance measurements

The e-pili conductance was measured using the same interdigitated electrode configuration as previously described (40). Briefly, 40 nm thick four-point probe gold electrodes were fabricated onto 300 nm thick silicon dioxide antimony-doped silicon wafers using photolithography. The nano-device was 4 mm by 4 mm with each of the gold pads measuring 1 mm by 1 mm. The interdigitated electrodes were 400 mm long with a non-equidistant gap of 15 µm between the 2 inner electrodes (SMU2 and SMU3) and 3 µm between the inner and outer electrodes (Source/SMU2 and SMU3/Drain). The electrodes were connected to a Keithley 4200 Semiconductor Characterization System Parametric Analyzer (Tektronix Inc., Beaverton, OR, USA) by 1 μm diameter tungsten probes (Signatone, Gilroy, CA, USA) via triaxial cables.

Each of the triplicate purified e-pili samples (2 μl, 500 μg protein/μl) were dropcast onto the interdigitated electrodes and allowed to air dry at 24 °C. After which, the current-voltage (I-V) curves for each of the triplicate devices was obtained in triplicate using a ±30 × 10−8 V sweep with a 5 second delay and a 250 second hold time. The conductance for each curve was calculated by extracting the slope of the linear fit of the current-voltage response for each of the technical replicates and subsequently for the biological replicates.

## Acknowledgements

We thank Professor Derek R. Lovley from the Department of Microbiology at the University of Massachusetts Amherst for reading the manuscript and providing helpful suggestions.

